# Adenovirus serotype 26 utilises sialic acid bearing glycans as a primary cell entry receptor

**DOI:** 10.1101/580076

**Authors:** Alexander T. Baker, Rosie Mundy, James Davies, Pierre J. Rizkallah, Alan L Parker

## Abstract

Adenoviruses are clinically important agents. They cause respiratory distress, gastroenteritis, and epidemic keratoconjunctivitis (EKC). As non-enveloped, double stranded DNA viruses, they are easily manipulated, making them popular vectors for therapeutic applications, including vaccines. Species D adenovirus serotype 26 (HAdV-D26) is both a cause of EKC and other disease, and a promising vaccine vector. HAdV-D26 derived vaccines are under investigation as protective platforms against HIV, Zika, RSV infections and are in Phase-III clinical trials for Ebola.

We recently demonstrated that HAdV-D26 does not utilise CD46 or desmoglein 2 as entry receptors, whilst the putative interaction with Coxsackie and Adenovirus Receptor (CAR) is low affinity and unlikely to represent the primary cell receptor.

Here, we definitively establish sialic acid as the primary entry receptor utilised by HAdV-D26. We demonstrate removal of cell surface sialic acid inhibits HAdV-D26 infection and provide a high-resolution crystal structure of HAdV-D26 fiber-knob in complex with sialic acid.

## Introduction

Adenoviruses are clinically significant, both as human pathogens, and as platforms for therapeutic applications. As pathogens, human adenoviruses (HAdV) have been isolated from severe infections in both immunocompromised and healthy individuals(*1, 2*). Some adenoviruses have been associated with acute infection of the eye(*3*), respiratory(*4, 5*), and gastrointestinal tract(*1, 6*). In rare cases infections prove fatal, as observed in a recent neonatal infection of HAdV-D56(*7*), among adult patients in New Jersey with HAdV-B7d(*8*), and infamously with HAdV-E4 where large epidemics of adenovirus infection have been seen in military recruits(*9, 10*). However, most fatal infections are observed in immunocompromised individuals(*11*), such as those suffering from graft versus host disease (GVHD)(*12*) or HIV(*13, 14*).

Adenoviruses are classified into 7 species (A-G)(*15*), and between 57 and 90 serotypes depending on the taxonomic definitions used(*16, 17*). Some adenoviruses have been studied in detail, having well defined receptor tropisms, including as coxsackie and adenovirus receptor (CAR)(*18, 19*), CD46 (Major Complement Protein, MCP)(*20–22*), desmoglein 2 (DSG2)(*23–25*), or sialic acid bearing glycans(*26–29*). However, most serotypes have low seroprevalence in the population(*30–33*), though this is varies significantly by geographical location(*34, 35*). Their rarity means many serotypes remain understudied, with poorly defined primary receptor interactions. This is especially true of the species D adenoviruses (HAdV-D); the largest of the adenoviral species, containing 35 of the 57 canonical serotypes(*17*).

Species D adenoviruses are associated with several pathogenicities. HAdV-D56 is a potentially fatal emergent respiratory pathogen comprised of a recombination between 4 species D adenoviruses(*7*). Opportunistic adenovirus infection isolated from HIV/AIDS patients are most commonly from species D, where they are associated with prolonged shedding in the gastrointestinal tract(*13*). HAdV-D has also been associated with genital disease(*36, 37*). The species D adenoviruses are best known, however, for causing epidemic keroconjunctivitis (EKC) infections, which is endemic, but not isolated, to Japan(*38, 39*). Classically the primary EKC causing adenoviruses have been HAdV-D8(*40–42*), 37(*42–44*), and 64(*42, 45*) (previously classified as 19a(*46*)). More recently other species D adenovirus have been associated with EKC, including HAdV-D53(previously classified as HAdV-D22/H8)(*47, 48*), 54(*49*), 56(*50*).

Their double stranded DNA genome makes them readily amenable to genetic modification(*51*), and therefore has made them attractive candidates for genetic manipulation for therapeutic applications in cancer (oncolytic viruses)(*52*) and as vaccine vectors(*53, 54*). Species D adenoviruses are of special interest as vaccine vectors. Their ability to induce robust cellular and humoral immunogenic responses in humans, coupled with low seroprevalence rates in the general population(*30, 33*) makes them attractive platforms for vaccines, as evidenced by their progression through clinical trials for HIV(*55, 56*), Zika(*57*), and Ebola treatment(*58, 59*). However, there remains a lack of understanding regarding their basic biology and mechanisms of cellular infection. This is exemplified by Adenovirus serotype 26 (HAdV-D26), which is being investigated as a vaccine vector for zika(*57*), HIV(*60*), respiratory syncytial virus(*61*), and has entered phase III clinical trials as an Ebola vaccine(*58*).

Despite its clinical success, recent findings further highlight the lack of clarity over the primary receptor usage of HAdV-D26. It is now clear that, despite previous publications to the contrary, HAdV-D26 cannot engage CD46 as a primary cellular entry receptor(*62*). Instead, the HAdV-D26 fiber-knob protein (HAdV-D26K) may engage CAR as a primary receptor, although the affinity of this interaction is attenuated compared to the classical HAdV-C5 interaction with CAR due to the presence of an extended HAdV-D26 fiber-knob DG loop, which sterically inhibits the interaction with CAR(*62*). The deduced low affinity of the interaction between CAR and HAdV-D26 fiber-knob make it unlikely that CAR represents the definitive primary receptor of HAdV-D26.

Here, we conclusively demonstrate that HAdV-D26 utilises sialic acid bearing glycans as a primary entry receptor, and that this interaction can form a productive infection. We deduce the structure of HAdV-D26K in complex with sialic acid (Neu5Ac), demonstrating a similar topology to the known sialic acid interacting adenovirus HAdV-D37 fiber-knob in the sialic acid binding pocket, but highlight crucial mechanistic differences likely to enhance HAdV-D26 affinity for sialic acid compared to other serotypes.

## Results

### HAdV-D26K has an electrostatic profile permissive to sialic acid interaction

Our recent findings rule out any role for DSG-2 or CD46 in HAdV-D26 infection, whilst the low affinity of the interaction between HAdV-D26K and CAR made it an unlikely primary cell entry receptor. Previous amino acid sequence alignments demonstrated little conservation of sialic acid binding residues with the fiber-knob domains of the known sialic acid utilising adenoviruses HAdV-G52SFK (short fiber-knob) or Canine adenovirus serotype 2 (CAV-2)(*62*). However, these alignments indicated HAdV-D37, known to bind sialic acid in the apex of the fiber-knob, bore some similarity at a sequence level. We sought to evaluate the ability of HAdV-D26K to interact with the remaining previously described adenovirus receptor, sialic acid.

HAdV-D37 fiber-knob is identical to that of HAdV-D64, and highly homologous to HAdV-D8 (Fig.1A). These three viruses have been shown to cause epidemic keratoconjunctivitis (EKC) and to interact with sialic acid. The closely related HAdV-D19p, differing from HAdV-D64 at only two residues, has also been shown to bind sialic acid, but does not cause EKC. We compared HAdV-D26K to these viruses to determine if a similar binding mechanism was possible.

**Figure 1:**
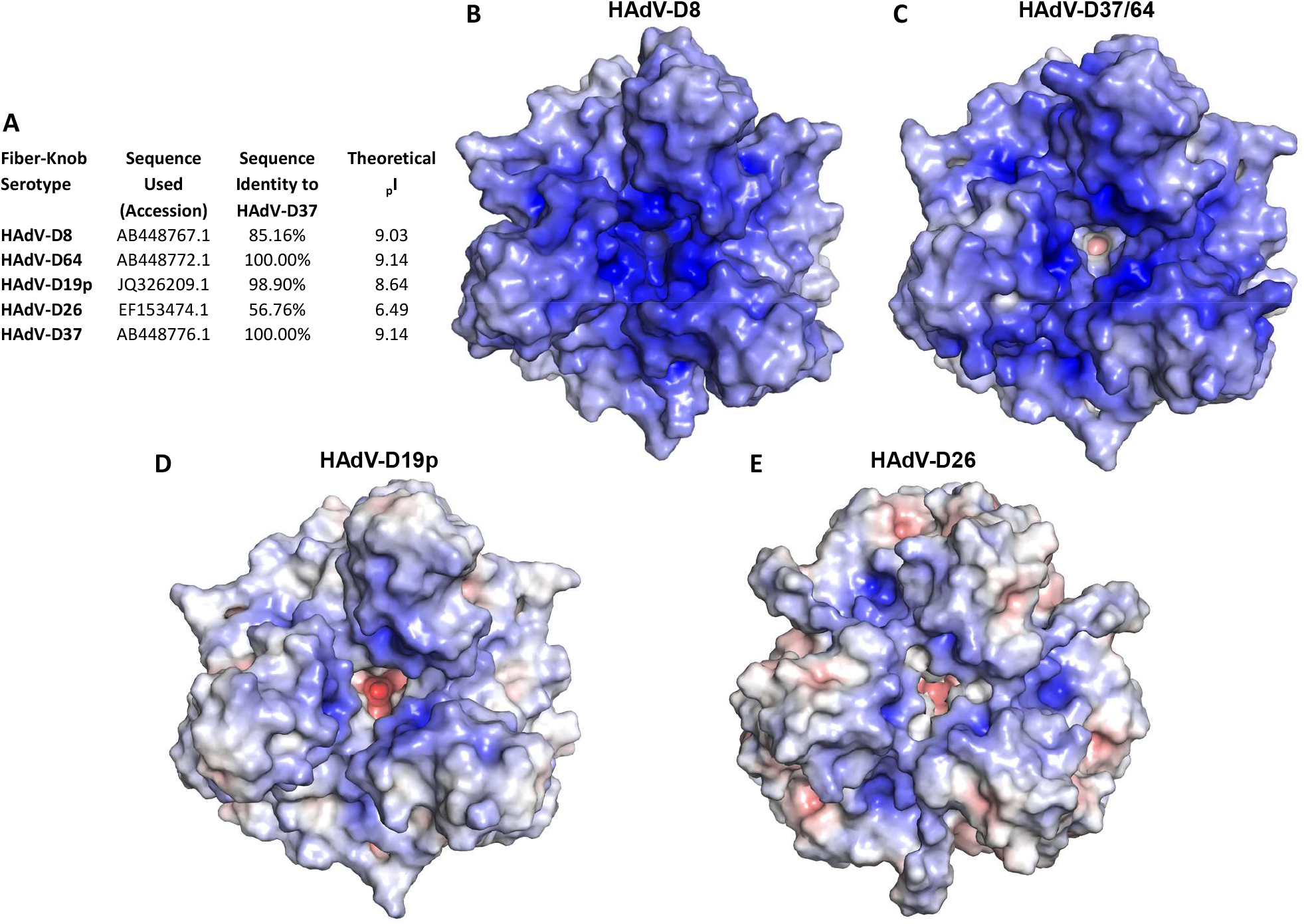
HAdV-D26K forms a local basic area in the apical depression to facilitate sialic acid binding despite net positive surface potential. HAdV-D26K has low (56.76%) sequence identity with fiber-knobs known to bind sialic acid by a similar mechanism, and an acidic isoelectric point (**A**). The electrostatic potential surfaces of HAdV-D8K (**B**), HAdV-D64/37 (**C**), and HAdV-D19p (**D**) fiber-knobs are highly basic, especially about the central depression about the 3-fold axis. HAdV-D26 fiber-knob is less basic overall but maintains positive potential in the central depression (**E**). Surfaces are displayed at ±10mV.

These sialic acid binding viruses all have highly negative predicted isoelectric points (pI) (Fig.1A). We calculated the surface electrostatic potentials of these fiber-knob proteins, at pH7.35 to simulate the pH of extracellular fluid, using previously published crystal structures where available. There is no published structure of HAdV-D8K, so we generated a homology model based on the closest known relative with a crystal structure (Fig.1B).

The viruses are highly basic, with a concentration of positive charge in the central depression around the 3-fold axis corresponding to the previously reported sialic acid binding sites (Fig. 1B-D). We observed that HAdV-D8 has the most basic surface potential (Fig.1B), followed by HAdV-D37/64 (Fig. 1C). HAdV-D19p is less basic, due to the two amino acid substitutions, compared to HAdV-D37/64, though the central depression is unaffected, as has previously been noted (Fig.1D)(*63*).

HAdV-D26K has a lower predicted pI, 6.49, and less positive surface potential (Fig.1A,E). However, the central depression of HAdV-D26K remains basic around the region where sialic acid is observed to bind in HAdV-D19p and HAdV-D37. HAdV-D26 retains the charge needed for sialic acid binding in the apex of the protein in the context of an otherwise acidic protein (Fig.1F).

### HAdV-D26 requires cell surface sialic acid for efficient infection

Sequence alignment of HAdV-D26K with these known sialic acid utilising viruses, bearing a positively charged apex, showed conservation of key binding residues between serotypes (Fig.2A). We observe complete conservation of Tyr130, and Lys165 across the 4 serotypes, and conservation of Asp128 with HAdV-D8 (Fig.2A). Further, while Tyr135 is not conserved in HAdV-D26, inspection of the crystal structure of HAdV-D37K and HAdV-D19p in complex with sialic acid (PDB 1UXA and 1UXB, respectively)(*63*) reveals this to be a main chain oxygen contact, positioned similarly in HAdV-D26, and can be considered homologous.

**Figure 2:**
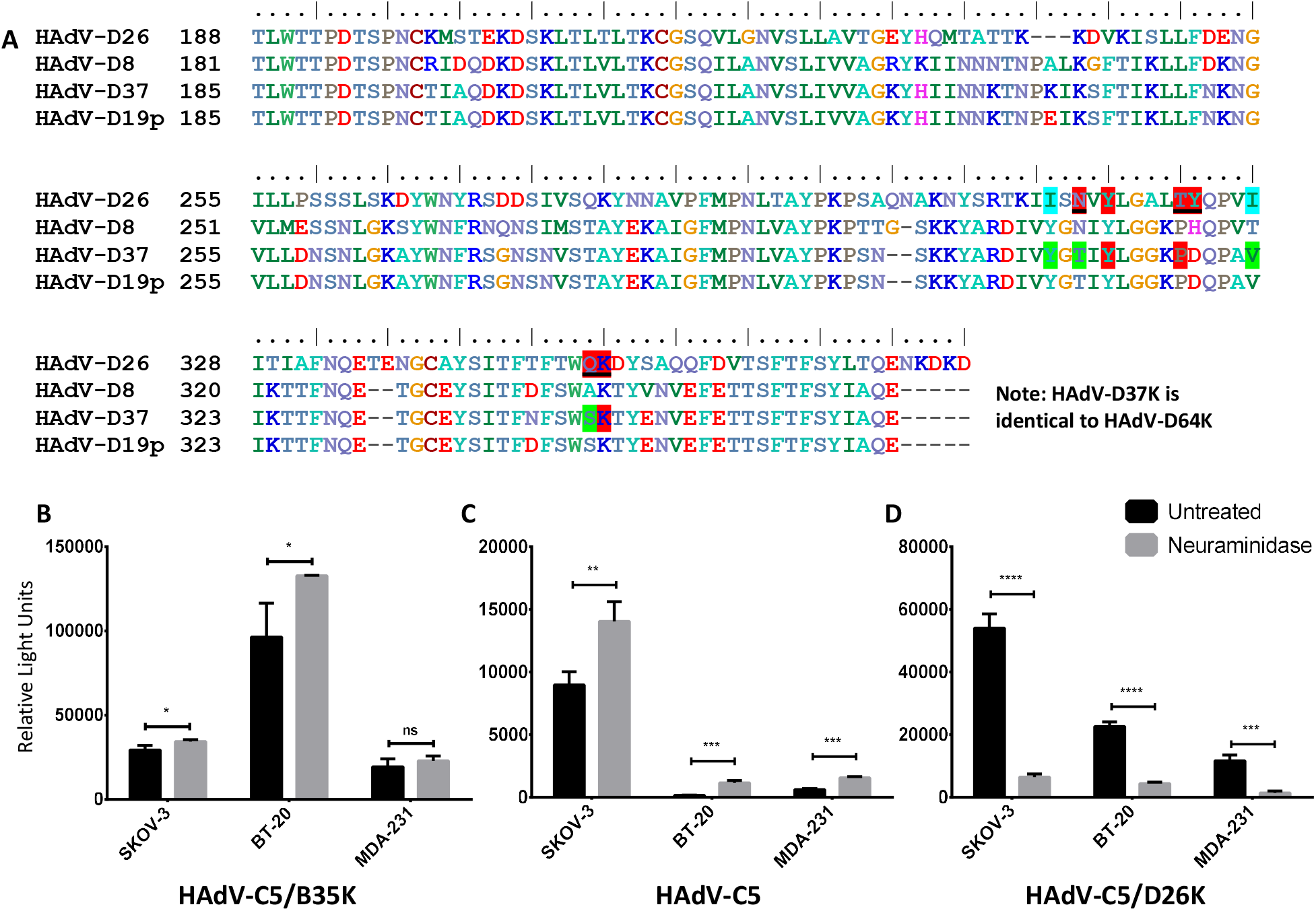
HAdV-D26K shares key binding residues with sialic acid utilising adenoviruses and exploits sialic acid to infect cells. Sequence alignment of HAdV-D26K shows conservation of key binding residues with known sialic acid binding adenoviruses (**A**). Residues highlighted in Red form polar contacts with sialic acid, green contact sialic acid via water-bridge, a black underline indicates both direct and water-bridge contacts, blue indicates hydrophobic contacts. Neuraminidase treatment does not reduce the ability of HAdV-D5/B35K (**B**) or HAdV-C5 (**C**) to infect SKOV-3 (ovarian adenocarcinoma), BT-20 (breast carcinoma), or MDA-231 (metastatic breast adenocarcinoma) cells, while HAdV-D5/D26K (**D**) is significantly inhibited. n=3 biological replicates, error = ±SD.

To investigate the ability of HAdV-D26 to utilise sialic acid as a cell entry receptor we used a replication incompetent HAdV-C5 vector pseudotyped with the HAdV-D26 fiber-knob, expressing a GFP transgene. We performed infectivity studies in three cell lines, with and without pre-treatment with neuraminidase to remove cell surface sialic acid. The tested cell lines could be infected by the CD46 (Fig.2B) or CAR (Fig.2C) mediated pathways to some extent, by HAdV-C5/B35K or HAdV-C5, respectively. However, infection via these routes was uninhibited by neuraminidase treatment. Transduction efficiency of HAdV-C5 and HAV-C5/B35K was actually enhanced by neuraminidase treatment in some cases; an effect which has been previously observed(*27, 64*). This has been suggested to be due to a reduction in the electrostatic repulsion of the negatively charged capsid of HAdV-C5.

Infection by the HAdV-C5/D26K pseudotype was significantly reduced in all three cell lines following treatment with neuraminidase (Fig.2D). This inhibition is significant (P<0.005), resulting in >5-fold decrease in infection, in all three cell lines tested. These data indicate that HAdV-C5/D26K is likely to be utilising the sialic acid mediated pathway for infection, not CD46 or CAR.

### HAdV-D26 forms a stable complex with sialic acid

We crystallised HAdV-D26K in complex with sialic acid in order to clarify the mechanism of interaction. Refinement of structures generated from HAdV-D26K crystals soaked in sialic acid shows electron density for a small molecule ligand in the apical depression (Fig.3A), this is best described by a racemic mixture of α and β anomers, in conjunction with double conformations of sialic acid (Fig.3B). The cubic space group (supplementary table 1) enabled assembly of the biological trimer. We observed three copies of sialic acid bound within the apex of the fiber-knob trimer (Fig.3C), as previously observed in HAdV-D37 and HAdV-D19p.

**Figure 3:**
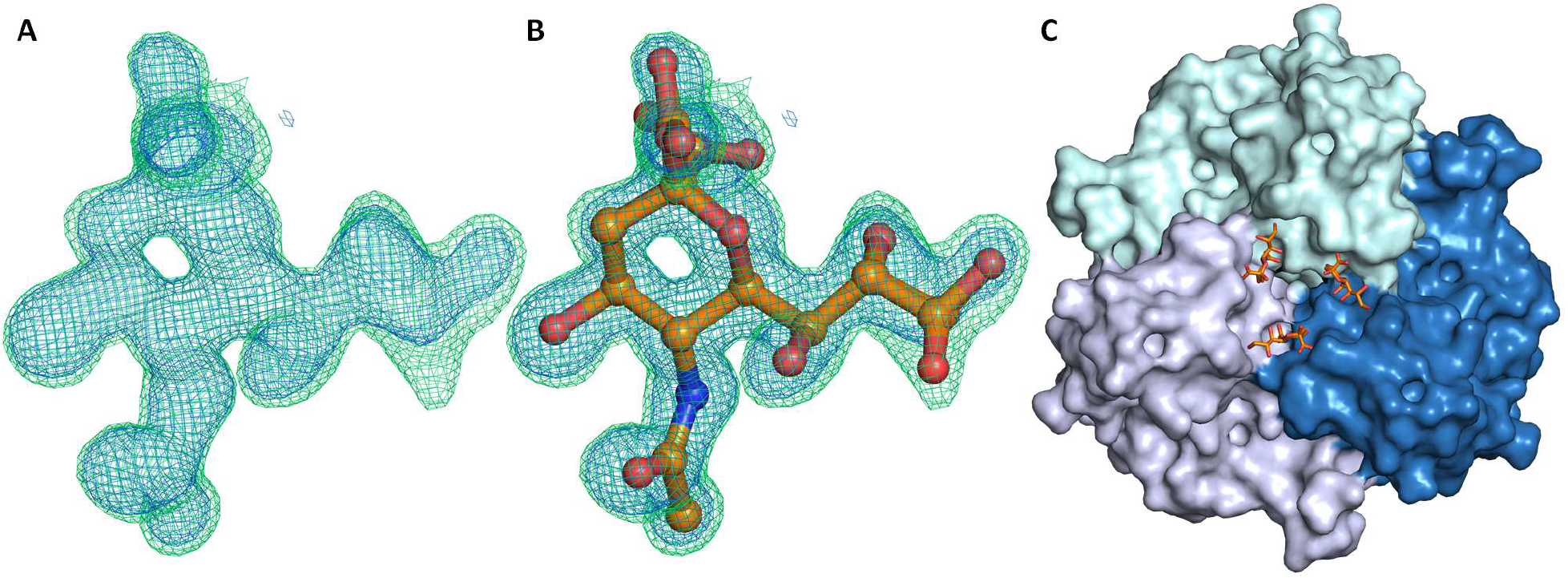
Sialic acid binds in the apical depression of adenovirus 26 fiber-knob protein. Sialic acid (orange) is seen to bind in 3 locations in the apical depression of the HAdV-D26 fiber-knob, bridging between monomers (shades of blue) of the trimeric assembly (**A**). The map shows clear density for a ligand (**B**), which is best described by a double conformer of sialic acid (**C**). Crystallisation statistics in supplementary table 1, 2FoFc map (blue mesh, σ=1.5), FoFc (Green mesh, σ=3.0).

Sialic acid binding was observed in structures crystallised at both pH8.0 (PDB 6QU6) and pH4.0 (PDB 6QU8). Observation of sialic acid density at high σ-values suggests a highly stable interaction (Supplementary Fig.1). Electron density demonstrates the C2 carboxyl and OH groups in two conformations, and the C6 glycerol group is flexible, with the C7-C8 bond rotating to alter the orientation of the glycerol arm relative to the pyranose ring and binding pocket (Fig.3A,B, Supplementary Fig.1). The glycerol group exhibits further flexibility at the C8-C9 bond, making the terminal oxygen mobile. The distribution of the density for the glycerol group is different at each pH (Supplementary Fig.1), suggesting pH could affect the preferred mode of interaction.

The most biologically relevant sialic acid conformation places the carboxyl group axial to the chair-conformation pyranose ring (Supplementary Fig.2), leaving the OH group pointing away from the fiber-knob and free to form an α(*2*)-glycosidic bond as part of a glycan. This is suggestive of a terminal sialic acid residue, as the chain can extend out of the central depression, as was observed in the previously described HAdV-D37K:GD1a glycan structure(*63, 65*).

### HAdV-D26 possesses a sophisticated sialic acid binding pocket

Comparison between the HAdV-D26K and HAdV-D37K, the best described of the sialic acid binding adenoviruses, reveals several sialic acid contacts are conserved (Fig.4A,B). Lys349 and Tyr314 are identical, and while Lys349 exhibits some flexibility, all observed lysine conformations form a contact with the carboxyl-group of the sialic acid (Supplementary Fig.3). Whilst Thr319 is not conserved in HAdV-D37 (which has a proline at this position), the main chain oxygen contact to the N-Acetyl nitrogen is spatially similar, so the bond can be considered homologous.

**Figure 4:**
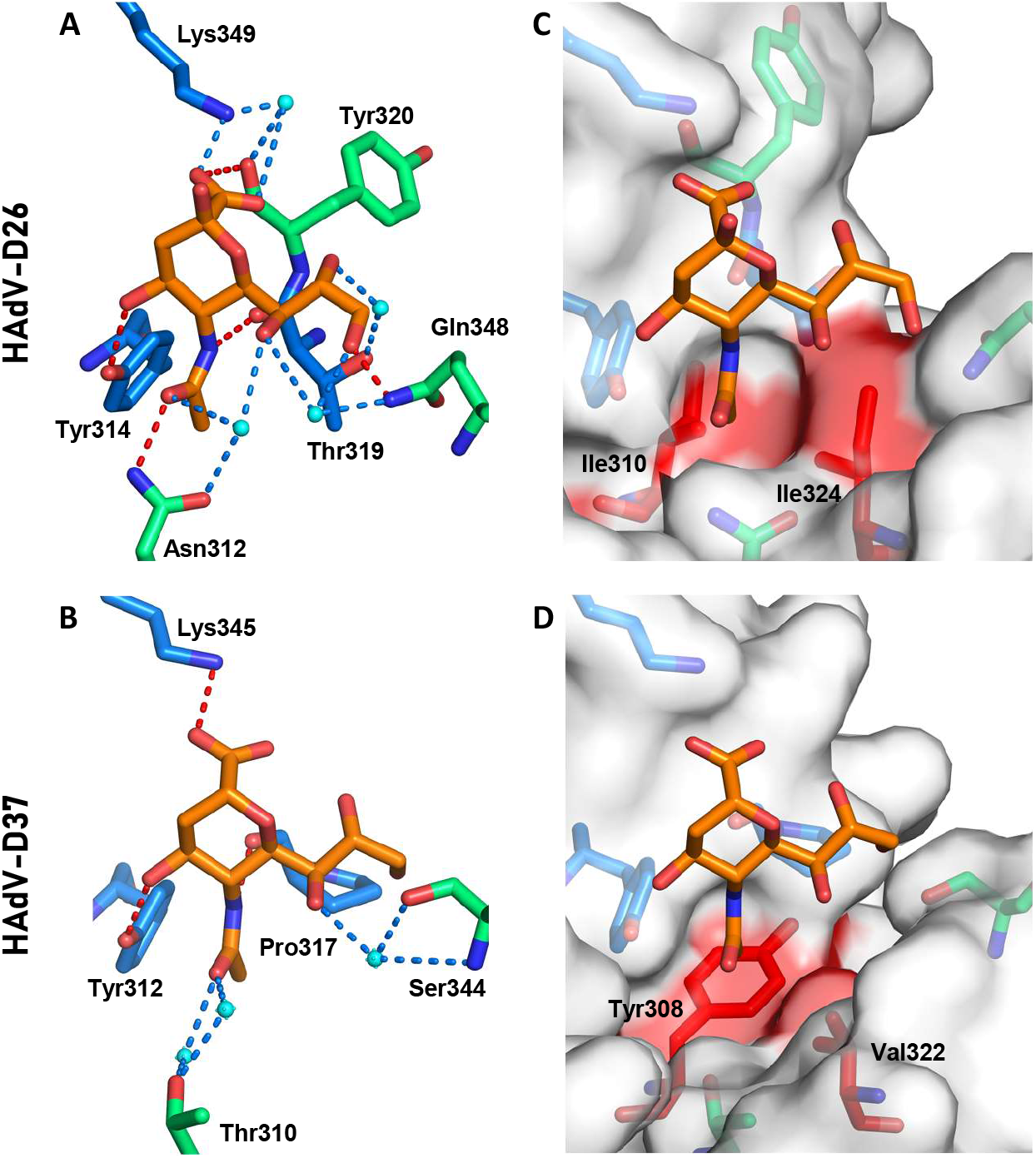
HAdV-D26K forms a complex interaction network of hydrophobic and electrostatic interactions with sialic acid. Sialic acid (orange) is seen to bind HAdV-D26 (**A**) and HAdV-D37 (**B**) through a network of polar contacts (red dashes) and hydrogen bonds (blue dashes). The interaction is stabilised by hydrophobic interactions (red regions on white surface) with the N-Acetyl CH3 group, but different residues in HAdV-D26 (**C**) and HAdV-D37 (**D**). Waters are shown as cyan spheres, residues forming comparable contacts in HAdV-D26 and HAdV-D37 are shown as blue sticks, other residues are shown as green sticks. Oxygen and nitrogen are seen in red and blue, respectively.

The HAdV-D26K sialic acid interface forms further contacts with sialic acid that are not observed in HAdV-D37K (Fig.4A, B). HAdV-D26K contacts the N-Acetyl oxygen of sialic acid using Asn312, which forms a polar contact and a water-bridge (Fig.4A). The comparable residue in HAdV-D37K, Thr310, is too short to form a direct polar interaction (Fig.4B), instead utilising a pair of water-bridges.

In HAdV-D37 the glycerol arm of sialic acid was only contacted by a water-bridge between Ser344 and the C7-OH. However, in HAdV-D26 all 3 OH groups in the glycerol arm form contacts. C7-OH is coordinated by water-bridges to both Asn312, and Gln348. C8-OH forms a water-bridge with Thr319, and C9-OH forms both a water-bridge and a polar contact directly to Gln348. Like Thr310, the serine belonging to HAdV-D37 at position 344 is too short to form a polar bond equivalent to the one with Gln348.

Notably, the density for the glycerol arm of sialic acid suggests several possible conformations (supplementary fig.2) which can be interpreted as flexibility. However, we suggest that, in HAdV-D26, this is unlikely since it is so well coordinated in all conformations observed, at both pH8.0 and pH4.0 (supplementary fig.3). We propose that HAdV-D26K can form a stable interaction with the glycerol arm, regardless of the specific confirmation. The variable density can be explained as the average distribution (or partition) of the different discrete positions.

We also observe a hydrophobic interaction in HAdV-D26 with the N-Acetyl methyl group at C11 (Fig.4C). A similar hydrophobic interaction is seen in HAdV-D37, where Tyr312 and Val322 form a hydrophobic patch (Fig.4D), but the HAdV-D37 interaction appears to be more selective, where Ile310, and Ile324 form a hydrophobic cradle around the methyl group (Fig.4C).

### HAdV-D26 binds sialic acid through an induced fit mechanism

We observe split density for Gln348 in both pH8.0 (Fig.5A) and pH4.0 (Fig.5B). Whilst conformation A can form polar contacts with sialic acid, conformation B points into the solvent and cannot. It is possible that Gln348 is flexible, but then is attracted to the charged density of the glycerol arm upon sialic acid interaction. We also observe greater occupancy of conformation A in the pH8.0 structure (approximately 0.7) while at pH4.0 the occupancy is evenly split. This suggests that the interaction may be more stable at higher pH, such as that associated with the pH found at the cell surface.

**Figure 5:**
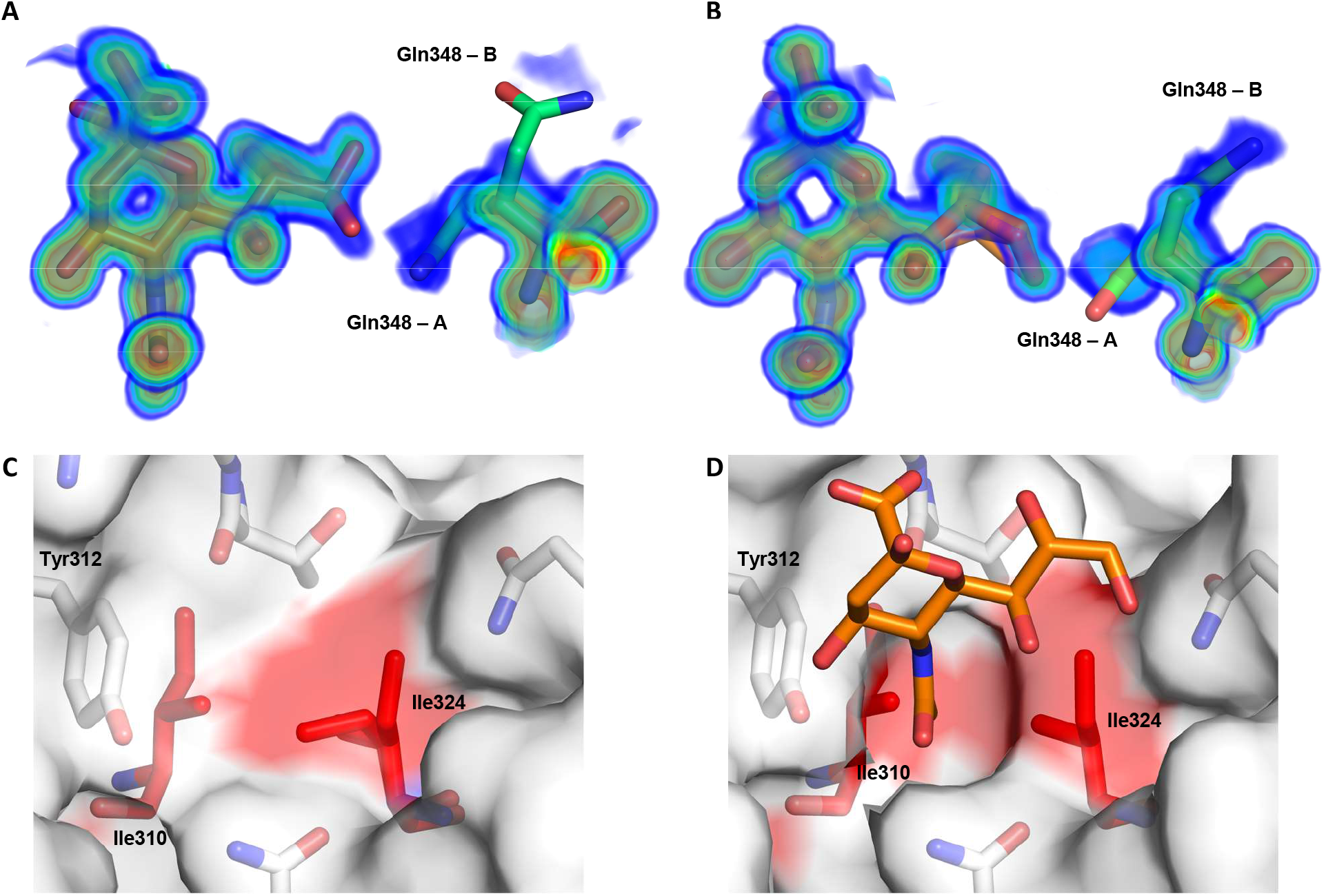
HAdV-D26K affects an induced fit mechanism in sialic acid binding. HAdV-D26K residue Gln348 can occupy multiple conformations, with a greater preference for conformation A (capable of forming a polar contact with the glycerol arm of sialic acid) at pH8.0 (**A**) than at pH4.0 (**B**). Ile324 has two conformations when HAdV-D26K is unliganded (**C**, PDB 6FJO). However, upon sialic acid binding the Ile324 adopts a single confirmation creating a hydrophobic indentation around the N-Acetyl methyl group bounded by Ile324,Ile310, and the ring of Tyr312 (**D**).

Ile324, which is seen to be involved in hydrophobic interactions with the N-Acetyl methyl group (Fig.4C), can also have multiple conformations. In an unliganded structure of HAdV-D26 fiber-knob (PDB 6FJO) the long arm of Ile324 is seen to rotate (Fig.5C). However, in the ligated structure Ile324 occupies a single conformation (Fig.5D) forming a cradle. This creates a larger hydrophobic patch and restricts the methyl group in space by pinching it between the pair of hydrophobic isoleucines, anchoring the N-Acetyl group.

## Discussion

Other adenoviruses have previously been shown to interact with sialic acid. These include CAV2(*66*), Turkey adenovirus 3 (TAdV-3)(*67*), and HAdV-G52 short fiber-knob(*26, 68*), but these viruses interact with sialic acid in lateral regions of the fiber-knob, dissimilar from HAdV-D26K. Four other human adenoviruses fiber-knob proteins (HAdV-D8/19p/37/64K) have been previously shown to utilise sialic acid, binding in the apical region. These viruses have high sequence similarity to each other, but not to HAdV-D26K, though they all share key sialic acid contact residues (Fig.1A,2A).

The structure of HAdV-D8 has not been determined, either alone or in complex with sialic acid, but infection by HAdV-D8 is sensitive to neuraminidase treatment suggesting sialic acid utilisation(*69*). Furthermore, HAdV-D8K has very high sequence homology and shared sialic acid contact residues with HAdV-D19p/37K making it logical to expect a similar interaction mechanism. In support of this we observe a similar electrostatic profile in the modelled fiber-knob as seen in HAdV-D37/64 (Fig.1B,C). HAdV-D64 has an identical fiber-knob domain to that of HAdV-D37, so fiber-knob interactions with sialic acid are likely to be conserved between these serotypes. HAdV-D26K conserves the key region of positive potential in the apical depression, but in the context of an otherwise more acidic protein (Fig.1E).

Inspection of the sialic binding pocket of HAdV-D26K reveals a much more complex mechanism of interaction than that previously reported for HAdV-D37K (Fig.4)(*63*). The overall topology of the pocket is similar, with hydrophobic residues around the N-Acetyl group and polar contacts between the carboxyl and C4-OH group. However, HAdV-D26K has several differences which increase the number of contacts between the sialic acid and the fiber-knob.

Subtle sequence changes enable more numerous interactions between HAdV-D26K and sialic acid than are possible in HAdV-D37K. In HAdV-D37 Pro317 forms a main chain oxygen contact to the nitrogen of sialic acid, however it also creates tension which rotates the N-terminal residue away from the carboxyl group of sialic acid. In HAdV-D26K, Tyr320, which is C-terminal of the Thr319 that is equivalent to Pro317 in HAdV-D37K does not create this tension and enables the main chain oxygen at position 320 to contact the sialic acid carboxyl group. Thr319, also forms a water-bridge with the C8-OH group, helping to stabilise the glycerol side-chain (Fig.4A).

This is one of several examples of HAdV-D26K being better evolved to contact sialic acid. The substitution of the Thr310 and Ser344 found in HAdV-D37K for longer charged residues (Asn312 and Gln348, respectively) in HAdV-D26K enables direct polar contacts, as well as additional water-bridge contacts. Substitution of Tyr308 and Val322 for more hydrophobic isoleucine residues in HAdV-D26 (Ile 310 and Ile324, respectively) creates a hydrophobic indentation better tailored to fit around the N-Acetyl methyl group.

The high resolution of the datasets generated to determine the sialic acid bound HAdV-D26K structure enables visualisation of multiple residue conformations with partial occupancy. In unligated structures of HAdV-D26K (PDB 6FJO), Ile324 exhibits a double conformer, occupying the available space (Fig.5C). However, when sialic acid is bound, it is restricted to have a single conformation with the long arm facing away from the sialic acid site, towards the inter-monomer cleft. Ile310 has the opposite orientation and creates an indentation which cradles the methyl group of sialic acid. Tyr312 may further contribute to the hydrophobic cradle. Tyr312 would not normally be considered a hydrophobic residue, but the side chain oxygen faces towards the solvent, where it forms a polar interaction with the C4-OH on sialic acid (Fig.4A), leaving the face of the tyrosine ring exposed to the methyl group which may contribute hydrophobic character to the cradle. This tyrosine behaves in both a polar, and hydrophobic manner at the same time. We suggest that the long arm of Ile324 adopts the sialic acid binding conformation in response to the hydrophobic pressure exerted by sialic acid entering the pocket, minimising the exposed hydrophobic surface when unbound, holding the methyl group between the short arm of Ile310 and the Tyr312 ring, making this an example of induced fit.

The double occupancy of Gln348 may indicate a second induced fit mechanism. We observe two possible conformations of Gln348 (Fig.5C). While conformation A does not form any contacts, conformation B forms a polar bond, and water-bridge, with the sialic acid glycerol group. In HAdV-D37K the glycerol group forms only a water-bridge from C7-OH to Ser344, the spatial equivalent of Gln348. Whilst we have not determined the preferred conformation of the sialic acid glycerol group versus Gln348 conformation, we observed that it can form polar contacts with it regardless, while Gln348 is in conformation B (Fig.5C).

We suggest Gln348 may be labile until the binding of sialic acid. Upon sialic acid binding Gln348 becomes attracted to the charged glycerol group causing it to stabilise in conformation A. This has the effect of “ locking” the glycerol side chain in place, which is further restrained by water-bridge contacts to Thr319 and Asn312.

Gln348 has greater occupancy in a sialic acid binding conformation (conformation-A) at pH8.0, which corresponds more closely to the physiological conditions in which it would encounter at the cell surface (Fig.5A). At pH4.0 Gln348 has approximately half occupancy in each conformation (Fig.5B). This implies the possibility of HAdV-D26K having lower sialic acid affinity under more acidic conditions, such as those encountered during endosomal trafficking down the lysosomal pathway.

Therefore, the HAdV-D26K binding pocket to sialic acid is summarised by three synchronous mechanisms. An N-Acetyl anchor comprised of a polar contact to Asn312 stabilised by a water bridge and an induced hydrophobic cradle around the methyl group. An inducible lock, where Gln348 forms a polar contact to the most terminal atoms in the glycerol arm, supported by a network of water bridges. Finally, a network of polar contacts to the carboxyl, C4 oxygen, and nitrogen atoms, which stabilise the pyranose ring.

This interaction in HAdV-D26K is a much more sophisticated binding mechanism compared to HAdV-D19p and the EKC causing viruses. However, the overall pocket topology and several key residues bear similarities. It may be surprising to observe such similarity given the low level of sequence homology HAdV-D26K has to the HAdV-D37K (56.76%, Fig.1A). Other regions, especially the loops, have highly dissimilar sequences. There is a precedent for this within adenovirus, with recombination events being reported in numerous settings(*46, 47, 70, 71*).

It has previously been suggested that many of the species D adenoviruses may have dual sialic acid binding affinity and CAR affinity(*63*). This has been observed in HAdV-D37/64, CAV-2(*19, 66*), and now HAdV-D26(*62*). Interestingly the species G adenovirus HAdV-G52 has also been observed to bind both CAR and sialic acid, but using two different fiber knob proteins on the same virus and a different mechanism of sialic acid interaction in the knob(*26*), which is shown to bind polysialic acid(*29*). Previous work has proposed CAR may be a receptor for many, if not all, of the species D adenoviruses with variable affinity(*62, 63, 72*), and suggest that sialic acid could also be widely utilised(*63*). These findings support that assertion, and adding another species D adenovirus, with low sequence similarity, to the pool of adenoviruses observed to bind both CAR and sialic acid.

Human adenovirus serotypes 43, 27, and 28 fiber-knobs share high sequence homology with HAdV-D26K, sharing the majority of the critical binding residues, and/or having structural homologues at those positions (Supplementary Fig.4). HAdV-D26K is the only species D adenovirus to have a glutamine at position 348 (HAdV-D26K numbering), though many have the shorter, but similarly charged, asparagine at this location, share the serine or similarly charged residue found in HAdV-D37K, or possess an asparagine which could behave similarly to glutamine. However, HAdV-D8 has an uncharged alanine at this position suggesting that a charged residue may not be strictly required for sialic acid binding, though may alter affinity (Supplementary Fig.4).

HAdV-D8K shares an asparagine at the same position as HAdV-D26K which we have shown to form polar and water-bridge contacts to sialic acid (Fig.4A). While this is unique among the classical EKC causing viruses HAdV-D19p/37/64, it is the most common residue at this position in the species D adenoviruses (Supplementary Fig.4).

The HAdV-D26K surface electrostatics are most like those of HAdV-D19pK. HAdV-D19pK is capable of binding sialic acid(*63*), and a limited effect is seen on infection of A549 cell binding after neuraminidase treatment to remove cell surface sialic acid(*69*). HAdV-D19p binding to Chang C (human conjunctival) cells was completely unaffected by neuraminidase treatment, though binding was very low regardless of neuraminidase treatment(*27*). This inability to bind Chang C cells was shown to depend upon a single lysine residue (Lys240) in the apex of the fiber-knob, but distant from the sialic binding pocket, creating a more acidic apical region in the lysine’s absence(*73*). HAdV-D26K also lacks a lysine in this position and has the most acidic electrostatic profile observed in this study (Fig.1).

HAdV-D37K, and the identical HAdV-D64K, have been shown to preferentially interact with the sialic acid bearing GD1a glycan on the corneal cell surface, causing EKC(*65*). However, it seems unlikely that a protein capable of trivalent sialic acid binding is completely specific for GD1a, a di-sialylated glycan, given the wide range of available glycan motifs which are di- and tri-sialylated. The GD1a preference may be diminished in HAdV-D19p by the acidic surface caused by the two amino acid substitutions, creating a glycan preference for tissues outside of the eye. Similarly, HAdV-D26K may have a unique glycan preference, driving its tissue tropism towards cells with different glycosylation patterns.

These findings clarify the receptor tropism of HAdV-D26 and build upon the increasingly complex body of knowledge describing species D adenoviruses. The comparison of different sialic acid binding residues suggests greater plasticity regarding the specific residues needed for sialic acid binding than previously thought (Supplementary Fig.4). It seems highly likely that many adenoviruses in species D, and perhaps other species, may interact with sialic acid in this manner. This suggests potential causes of off target infection by species D derived viral vectors. Conversely, investigation of their specific glycan preferences may enable more tissue specific targeting. Knowledge of the sialic acid binding mechanism suggests mutations which may ablate sialic acid interaction, enabling engineering of better restricted tropisms for future virotherapies. This knowledge regarding HAdV-D26 receptor can inform clinical practice in the rare cases of acute HAdV-D26 infection, or in the face of adverse reactions to HAdV-D26 based vaccines, suggesting that sialic acid binding inhibitors, such as Zanamivir, or trivalent sialic acid derivatives(*74*) may make effective anti-HAdV-D26 therapies.

## Materials and Methods

### Infectivity assays

Cells were seeded at a density of 30,000 cells/well in a flat bottomed 96 well cell culture plate and incubated overnight at 37°C to adhere. Cells were washed twice with 200μl of PBS and 50ul of neuraminidase (Sigma-Aldrich, Cat#11080725001) was added to the appropriate wells at a concentration of 50mU/ml, diluted in serum free media, and incubated for 1hr at 37°C. Cells were cooled on ice and washed twice with 200μl of PBS. Green Fluorescent Protein (GFP) expressing, replication incompetent viruses were added to the appropriate wells at a concentration of 2000 or 5000 viral particles per cell, in 100ul of serum free media, at 4°C, and incubated on ice for 1hr. Serum free media alone was added to uninfected control wells. Cells were washed twice with 200μl of cold PBS, complete media added (DMEM, 10% FCS) and incubated for 48hrs at 37°C. Cells were then trypsinised and transferred to a 96 well V-bottom plate, washed twice in 200μl of PBS and fixed in 2% paraformaldehyde for 20mins before wash, and resuspension in 200μl of PBS.

Samples were run in triplicate and analysed by flow cytometry on Attune NxT (ThermoFisher), analysed using FlowJo v10 (FlowJo, LLC), gating sequentially on singlets, cell population, and GFP positive cells. Levels of infection were defined as the percentage of GFP positive cells (%+ve), and/or Total Fluorescence (TF), defined as the percentage of GFP positive cells multiplied by the median fluorescent intensity (MFI) of the GFP positive population. These measures are distinct in that %+ve describes the total proportion of cells infected, and TF describes the total efficiency of transgene delivery.

### Amino Acid Sequence Alignments

Representative whole genomes of HAdV-D64, HAdV-D19p, HAdV-D26, and HAdV-D37 were selected from the National Center for Biotechnology Information (NCBI), the fiber-knob domain amino acid sequences were derived from them, defined as the translated nucleotide sequence of the fiber protein (pIV) from the conserved TLW hinge motif to the protein C-terminus. The fiber-knob domains were aligned using the EMBL-EBI Clustal Omega tool(*75*).

### Generation of Recombinant Fiber-Knob protein

SG13009 *E.coli* harbouring pREP-4 plasmid and pQE-30 expression vector containing the fiber-knob DNA sequence were cultured in 20ml LB broth with 100μg/ml ampicillin and 50μg/ml kanamycin overnight from glycerol stocks made in previous studies(*76–78*). 1L of TB (Terrific Broth, modified, Sigma-Aldrich) containing 100μg/ml ampicillin and 50μg/ml were inoculated with the overnight *E.coli* culture and incubated at 37°C until they reached OD0.6. IPTG was then added to a final concentration of 0.5mM and the culture incubated at 37°C for 4hrs. Cells were then harvested by centrifugation at 3000g, resuspended in lysis buffer (50mM Tris, pH8.0, 300mM NaCl, 1% (v/v) NP40, 1mg/ml Lysozyme, 1mM β-mercaptoethanol), and incubated at room temperature for 30mins. Lysate was clarified by centrifugation at 30,000g for 30mins and filtered through a 0.22μm syringe filter (Millipore, Abingdon, UK). Clarified lysate was then loaded onto a 5ml HisTrap FF nickel affinity chromatography column (GE) at 2.0ml/min and washed with 5 column volumes into elution buffer A (50mM Tris [pH8.0], 300mM NaCl, 1mM β-mercaptoethanol). Protein was eluted by 30min gradient elution from buffer A to B (buffer A + 400mM Imidazole). Fractions were analysed by reducing SDS-PAGE, and Fiber-knob containing fractions further purified using a superdex 200 10/300 size exclusion chromatography column (GE) in crystallisation buffer (10 mM Tris [pH 8.0] and 30 mM NaCl). Fractions were analysed by SDS-PAGE and pure fractions concentrated by centrifugation in Vivaspin 10,000 MWCO (Sartorius, Goettingen, Germany) proceeding crystallisation.

### Crystallisation and structure determination

Protein samples were purified into crystallisation buffer (10 mM TRIS [pH 8.0] and 30 mM NaCl). The final protein concentration was approximately 10 mg/ml. Commercial crystallisation screen solutions were dispensed into 96-well plates using an Art-Robbins Instruments Griffon dispensing robot (Alpha Biotech, Ltd), in sitting-drop vapour-diffusion format. Drops containing 200nl of screen solution and 200nl of protein solution were equilibrated against a reservoir of 60μl crystallisation solution. The plates were sealed and incubated at 18°C.

Crystals of HAdV-D26K appeared in PACT Premier condition B01 and B04 (0.1 M MIB [Malonic acid, Imidazole, Boric acid], 25 % w/v PEG 1500, pH4.0 and pH8.0 respectively), within 1 to 7 days. Crystals were then soaked in reservoir solution containing N-Acetylneuraminic acid (Neu5Ac, Sigma-Aldrich Cat#A2388) at a final concentration of 10mM and incubated overnight prior to harvest. Crystals were cryoprotected with reservoir solution to which ethylene glycol was added at a final concentration of 25%. Crystals were harvested in thin plastic loops and stored in liquid nitrogen for transfer to the synchrotron. Data were collected at Diamond Light Source beamline I04, running at a wavelength of 0.9795Å. During data collection, crystals were maintained in a cold air stream at 100°K. Dectris Pilatus 6M detectors recorded the diffraction patterns, which were analysed and reduced with XDS(*79*), Xia2, DIALS(*80*), and Autoproc(*81*). Scaling and merging data was completed with Pointless, Aimless and Truncate from the CCP4 package(*82*). Structures were solved with PHASER, COOT was used to correct the sequences and adjust the models, REFMAC5 was used to refine the structures and calculate maps. Graphical representations were prepared with PyMOL(*83*). Reflection data and final models were deposited in the PDB database with accession codes: 6QU6, 6QU8, and 6FJO. Full crystallographic refinement statistics are given in Supplementary Table 1

### Calculation of electrostatic surface potentials and isoelectric points

HAdV-D37, HAdV-D19p, and HAdV-D26 used PDB IUXA, PDB 1UXB, and PDB 6QU8, respectively, as the input. HAdV-D8 was calculated using a homology model, generated as described below, for input.

The PDB2PQR server (V 2.1.1)(*84*) was used assign charge and radius parameters using the PARSE forcefield, and assigned protonation states using PROPKA, at pH7.35. APBS(*85*) was used to calculate electrostatic surface potentials, and the map output was visualised in PyMol(*83*).

### Homology modelling of Adenovirus serotype 8

The I-TASSER protein structure and function prediction server(*86–88*) was used to generate a homology model of HAdV-D8 based on the published sequence of HAdV-D8(*42*), using the published structure of it’s closest relative (by sequence identity), HAdV-D19p(*63*). The resultant monomer was then copied three times, using the HAdV-19p trimer as a template, and the monomers aligned in PyMol(*83*) so as to generate a model of the complete HAdV-D8K trimer.

**Table 1.** Data collection and refinement statistics for structures generated in this study

| PDB Entry | 6QU6 | 6QU8 | 6FJO |
| --- | --- | --- | --- |
| **Data Collection** |  |  |  |
| Diamond Beamline | I04 | I04 | I04 |
| Date | 25/10/2018 | 25/10/2018 | 05/12/2017 |
| Wavelength | 0.91587 | 0.91587 | 0.9795 |
| **Crystal Data** |  |  |  |
| Crystallisation Conditions | 0.1 M MIB, 25 % w/v PEG 1500 | 0.1 M MIB, 25 % w/v PEG 1500 | 0.1 M SPG, 25% PEG 1500 |
| pH | 4.0 | 8.0 | 4.0 |
| $a=b=c$ (Å) | 85.73 | 85.92 | 85.78 |
| $\alpha=\beta=\gamma$ (°) | 90.0 | 90.0 | 90.0 |
| Space group | P 2<sub>1</sub> 3 | P 2<sub>1</sub> 3 | P 2<sub>1</sub> 3 |
| Resolution (Å) | 1.03 – 49.5 | 1.19 – 42.96 | 1.17-85.78 |
| Outer shell | 1.03 – 1.06 | 1.19 – 1.22 | 1.17-1.23 |
| *R*-merge (%) | 5.3 (125.3) | 8.5 (276.1) | 6.5 (137.0) |
| *R*-meas (%) | 5.5 (161.3) | 8.8 (283.2) | 6.6 (140.4) |
| CC1/2 | 1.0 (0.224) | 1.0 (0.505) | 1.00 (0.825) |
| *I* / $\sigma$(*I*) | 21.9 (0.7) | 20.7 (1.3) | 24.8 (2.3) |
| Completeness (%) | 97.7 (76.5) | 100.0 (100.0) | 100.0 (100.0) |
| Multiplicity | 14.9 (2.0) | 21.4 (20.3) | 21.8 (21.2) |
| Total Measurements | 1,509,159 | 1,448,478 | 1,568,641 |
| Unique Reflections | 103,975 | 67,799 | 71,878 |
| Wilson B-factor(Å<sup>2</sup>) | 8.4 | 11.1 | 12.2 |
| **Refinement Statistics** |  |  |  |
| Total number of refined | 1,936 | 1,846 | 1,761 |
| R-work reflections | 96,027 | 64,286 | 68,283 |
| R-free reflections | 4,907 | 3,441 | 3,558 |
| R-work/R-free (%) | 13.6 / 14.8 | 14.17 / 17.20 | 17.0 / 19.0 |
| **rms deviations** |  |  |  |
| Bond lengths (Å) | 0.012 | 0.011 | 0.021 |
| Bond Angles (°) | 1.754 | 1.661 | 2.080 |
| <sup>1</sup>Coordinate error | NULL | NULL | 0.026 |
| Mean B value (Å<sup>2</sup>) | 17.6 | 29.6 | 19.9 |
| **Ramachandran Statistics** |  |  |  |
| Favoured/allowed/Outliers | 119 / 9 / 0 | 126 / 10 / 0 | 133 / 10 / 1 |
| % | 93.0 / 7.0 / 0.0 | 92.7 / 7.4 / 0.0 | 92.4 / 6.9 / 0.7 |

\* One crystal was used for determining each structure.
\* Figures in brackets refer to outer resolution shell, where applicable.
<sup>1</sup> Coordinate Estimated Standard Uncertainty in (Å), calculated based on maximum likelihood statistics.
Buffers:
- MIB: Malonic acid, Imidazole, Boric acid - SPG: Succinic acid, Sodium phosphate monobasic monohydrate, Glycine: pH 4.0

## Acknowledgements

A.T.B is supported by a Tenovus Cancer Care PhD studentship to A.L.P (project reference PhD2015/L13). J. A. D is supported by a Cancer Research UK Biotherapeutics Drug Discovery Project Award to A.L.P. (project reference C52915/A23946). A.L.P and P.R are funded by Higher Education Funding Council for Wales.

The authors wish to acknowledge the Diamond Light Source for beamtime (proposal mx14843), and the staff of beamline I04 for assistance with diffraction data collection.

---

**Supplementary Figure 1:**
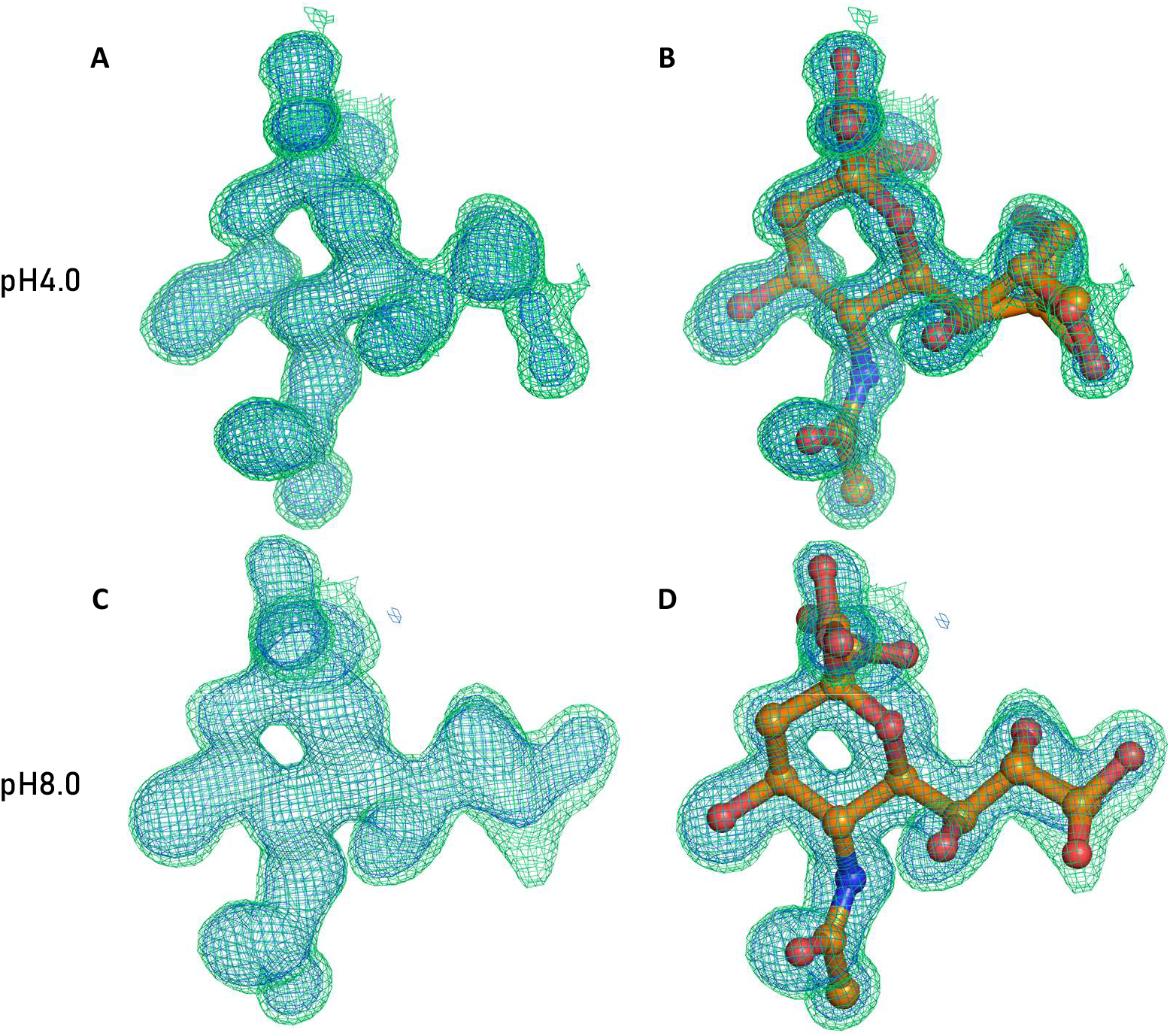
Sialic acid forms a stable interaction with HAdV-D26K at both pH4.0 (PDB 6QU6) and pH8.0 (PDB 6QU8). The omit map at pH4.0 (**A**) shows density for a small molecule ligand, which can be best modelled by a sialic acid double conformer (**B**). The same is true at pH8.0 (**C**), but the preferred conformations of the glycerol group are different (**D**). Crystallisation statistics in supplementary table 1, 2FoFc map (blue mesh, σ=1.5), FoFc (Green mesh, σ=3.0).

**Supplementary Figure 2:**
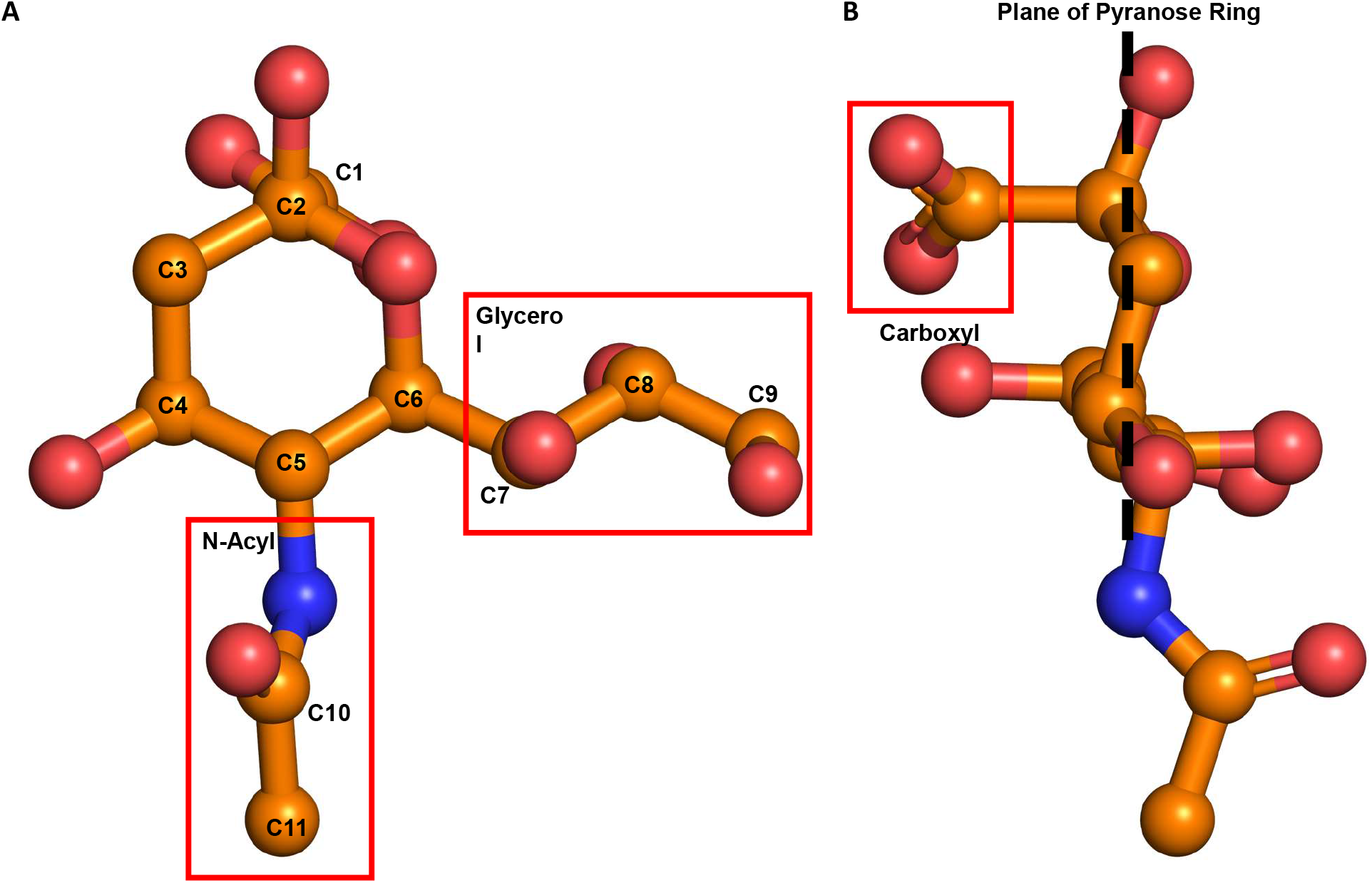
Structure of sialic acid (N-Acetylneuraminic acid, Neu5Ac) in a biologically relevant conformation. Viewing the Neu5Ac face on with glycerol and N-Acetyl groups labelled in red boxes, and the carbons numbered (**A**). Side on the carboxyl group (red box) is seen in the axial conformation with the C2 OH group planar to the ring (**B**).

**Supplementary Figure 3:**
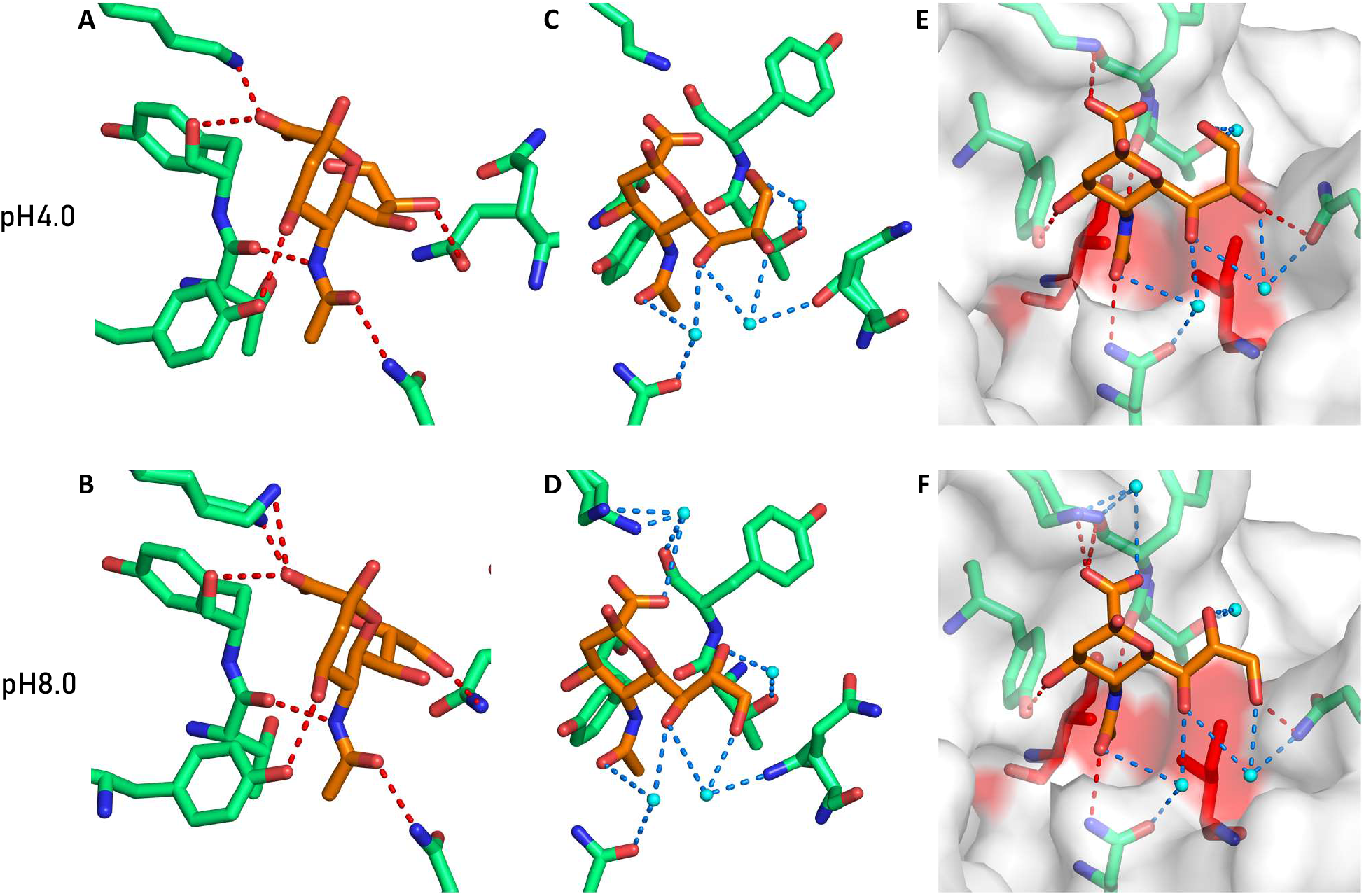
HAdV-D26K forms a similar interaction with sialic acid at both pH4.0 (PDB 6QU6) and pH8.0 (PDB 6QU8) through a combination of polar, water-bridge, and hydrophobic interactions. At pH4.0 sialic acid forms numerous polar contacts to charged side chains in HAdV-D26K (**A**), similar contacts are seen at pH8.0 and exhibits a lysine double conformer (**B**). At pH4.0 sialic acid forms several water bridges stabilising the interaction of the glycerol group (**C**), the same bridges are seen at pH8.0 with the addition of a water-bridge contact on the carboxyl group not seen at pH4.0 (**D**). A hydrophobic interface is formed around the N-Acetyl methyl group which appears to be similar and stable at pH4.0 (**E**) and pH8.0 (**F**). Polar bonds to residues are shown as red dashes, water-bridge contacts as blue dashes, and waters as cyan spheres. Sialic acid is shown in orange, polar HAdV-D26K residues as green sticks, and purely hydrophobic residues as red sticks. The HAdV-D26 surface is shown in white with hydrophobic regions in red. Oxygen and nitrogen atoms are coloured red and blue, respectively.

**Supplementary Figure 4:**
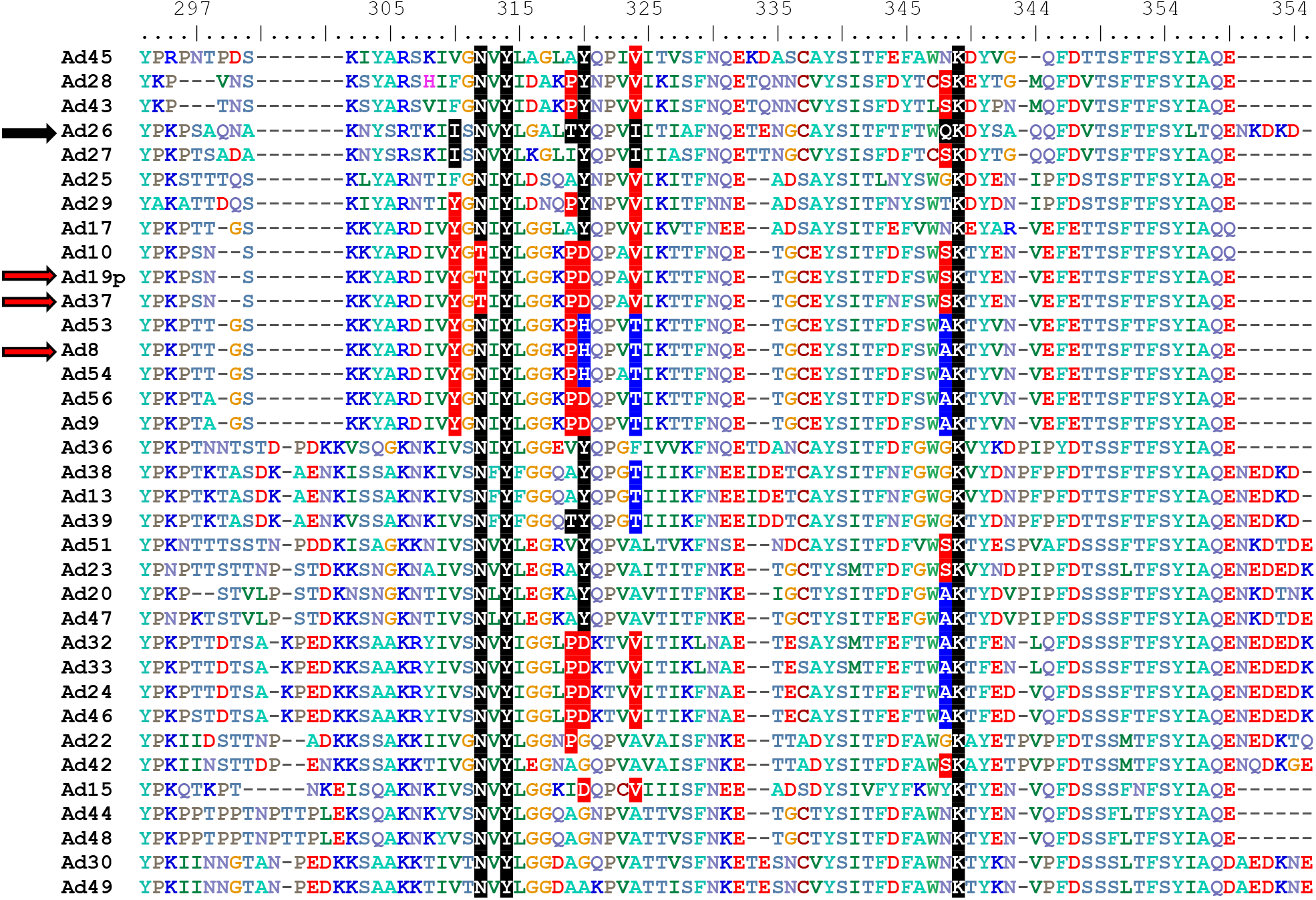
Species D adenoviruses conserve known sialic acid binding residues. Sequence alignment of species D adenovirus fiber-knob proteins. Known HAdV-D26 and HAdV-D37/64/19p residues forming contacts with sialic acid are highlighted in black and red respectively. Homologous residues are coloured similarly to the virus which they share the residue with. HAdV-D8 residues at known sialic acid binding locations which are dissimilar to HAdV-D26/37 are highlighted in blue, as are homologous residues in other viruses. Names utilise the short nomenclature, all are human species D adenoviruses. Numbering is for HAdV-D26K.

